# Loss of long-chain acyl-CoA dehydrogenase protects against acute kidney injury

**DOI:** 10.1101/2024.10.22.619640

**Authors:** Takuto Chiba, Akira Oda, Yuxun Zhang, Joanna Bons, Sivakama S Bharathi, Katherine E. Pfister, Bob B. Zhang, Adam C. Richert, Birgit Schilling, Eric S. Goetzman, Sunder Sims-Lucas

## Abstract

Proximal tubular epithelial cells (PTECs) are particularly vulnerable to acute kidney injury (AKI). While fatty acids are the preferred energy source for PTECs via fatty acid oxidation (FAO), FAO-mediated H_2_O_2_ production in mitochondria has been shown to be a major source of oxidative stress. We have previously shown that a mitochondrial flavoprotein, long-chain acyl-CoA dehydrogenase (LCAD), which catalyzes a key step in mitochondrial FAO, directly produces H_2_O_2_ *in vitro*. Further we have established that loss of a lysine deacylase, Sirtuin 5 (*Sirt5^−/−^*), induces hypersuccinylation and inhibition of mitochondrial FAO genes to stimulate peroxisomal FAO and to protect against AKI. However, the role of LCAD has yet to be determined.

Mass spectrometry data acquisition revealed that LCAD is hypersuccinylated in *Sirt5^−/−^* kidneys after AKI. Following two distinct models of AKI, cisplatin treatment or renal ischemia/reperfusion (IRI), LCAD knockout mice (*LCAD^−/−^*) demonstrated renoprotection against AKI. Specifically, *LCAD^−/−^* kidneys displayed mitigated renal tubular injury, decreased oxidative stress, preserved mitochondrial function, enhanced peroxisomal FAO, and decreased ferroptotic cell death.

LCAD deficiency confers protection against two distinct models of AKI. This suggests a therapeutically attractive mechanism whereby preserved mitochondrial respiration as well as enhanced peroxisomal FAO by loss of LCAD mediates renoprotection against AKI.

## INTRODUCTION

Dysregulated energy metabolism has long been recognized as a fundamental element of AKI, yet no effective therapeutic modalities have been developed to specifically target renal metabolic dysfunction^1^. Mitochondrial FAO, the process by which carbon chains of fatty acids are shortened for energy production, has been implicated as a source of H_2_O_2_ and oxidative stress in various tissues including kidneys^2,3^. PTECs, which are the main site of damage during AKI, are rich in mitochondria and rely on FAO for their high energy demands^4,5^. Previous studies in AKI have shown that mitochondrial reactive oxygen species (ROS) build up is a pivotal driver of cellular injury and cell death^6–8^.

LCAD catalyzes the first step in mitochondrial FAO. Mitochondrial FAO is integrated with the electron transport chain that produces ROS^9^; We recently showed that LCAD in the presence of fatty acids can lead to the direct production of H_2_O_2_ as a biproduct *in vitro*^10^. In addition, LCAD is downregulated in many cancers, possibly as a mechanism to reduce H_2_O_2_ levels^11,12^. Lysine acylation is a known posttranslational modification to inhibit LCAD activity^13,14^. It is not clear how LCAD is regulated during AKI or how it may contribute to tissue injury or recovery.

Here, we show via mass spectrometry data analysis that loss of the mitochondrial lysine deacylase Sirt5 caused LCAD hypersuccinylation which inhibits LCAD activity. We demonstrate that loss of LCAD in mice confers renoprotection against cisplatin and renal IRI, which are two distinct mouse models of AKI. Renoprotection in these models involves preservation of mitochondrial respiration and enhanced peroxisomal FAO. *LCAD^−/−^*mitigates oxidative stress and ferroptosis (oxidative stress induced cell death) after AKI. These results provide evidence for a potential novel therapeutic approach for the treatment of AKI.

## METHODS

### Succinylome analysis of *Sirt5^-/-^* and WT kidney tissues

We have previously reported the quantitative analysis of protein succinylation in *Sirt5^−/−^* and WT (n = 3) mouse kidney tissues collected after renal IRI^15^. The data is publicly available at: Mass Spectrometry Interactive Virtual Environment (MassIVE; ftp://massive.ucsd.edu/v02/MSV000083439/) with MassIVE ID: MSV000083439; ProteomeXchange ID: PXD012696. Briefly, kidney tissues were homogenized, trypsinized, desalted, and further enriched for succinylated peptides using the PTMScan Succinyl-Lysine Motif Kit (Cell Signaling Technologies) for succinyl-lysine PTM analysis. Samples were analyzed by data-independent acquisition on a TripleTOF 6600 system. The peptide AFG**^322^Ksucc**TVAHIQTVQHK with a succinyl group on the lysine residue K322 of the mitochondrial LCAD was analyzed in Skyline^16^. The extracted ion chromatograms of the peptide AFG**^322^Ksucc**TVAHIQTVQHK (precursor ion at m/z 441.99, z = 4+) were manually assessed and we confirmed accurate peak integration boundaries and removed potentially interfered transitions. Quantification and statistics were performed with Skyline: briefly peptide quantification was obtained by summing the transition area, and a *t*-test was applied to compare AFG**^322^Ksucc**TVAHIQTVQHK peptide abundance level in *Sirt5^−/−^* vs WT replicates.

### Animals

All animal protocols were approved by the University of Pittsburgh Institutional Animal Care and Use Committee (IACUC, protocol #22112009), and all experiments were conducted in accordance with the guidelines and regulations set forth in the Animal Welfare Act (AWA) and PHS Policy on Humane Care and Use of Laboratory Animals. *LCAD^−/−^* mice were obtained from the Mutant Mouse Regional Resource Center. The *LCAD^−/−^* mice on a mixed background of C57Bl/6 and 129 were crossed with a C57Bl/6 to generate heterozygous *LCAD^+/−^* mutants. Homozygous *LCAD^−/−^* mutants or WT controls (*LCAD^+/+^*) were derived from the common breeding pairs of *LCAD^+/−^*heterozygous mutants and used throughout the current study.

### Renal IRI induced AKI Model

Renal IRI was induced for the male mice by a unilateral renal IRI with a delayed contralateral nephrectomy as previously described^15,17,18^. Briefly, mice were anesthetized with 2% isoflurane. Core body temperature of the mice was monitored with a rectal thermometer probe and was maintained at 36.8-37.2°C throughout the procedures with a water-heating circulation pump system (EZ-7150; BrainTree Scientific) and an infrared heat lamp (Shat-R-Shield). Extended-release buprenorphine 1.3 mg/mL (Ethiqa XR, Fidelis MIF 900-014) was administered for analgesia (3.25 mg/kg b.w, s.c.). With aseptic techniques, a dorsal incision was made to expose the left kidney, and renal ischemia for 18 minutes was induced by clamping of the left kidney pedicle with a nontraumatic micro serrefines (18055–04; Fine Science Tools). Renal reperfusion was visually verified afterwards. Contralateral nephrectomy of the right kidney was performed at day 6 after renal IRI. Serum and the injured left kidney were collected at day 7. Serum was analyzed by the Kansas State Veterinary Diagnostic Laboratory for *Renal Profile* which include analysis of albumin, BUN, creatinine, and phosphorus.

### Cisplatin nephrotoxic AKI Model

To induce cisplatin AKI, male and female mice were given a single dose of cisplatin (20 mg/kg b.w. i.p. for females and 24 mg/kg b.w. i.p. for males; Fresenius Kabi), or vehicle control of normal saline as described previously^15, 18^. Serum and kidneys were collected on day 3 post-cisplatin.

### Sex as a biological variable

Both male and female mice were studied in the cisplatin AKI model. Male mice were used for the renal IRI AKI model.

### Western blotting

Kidney tissues were lysed in RIPA buffer (Thermo Fisher Scientific), and the homogenates plated in triplicate to measure protein concentration using a Bradford assay kit (Bio-Rad). Western blotting analysis was performed as previously described^15^. The following commercially available antibodies were used in this study: anti-GPX4 (Abcam ab125066), anti-PEX5 (Thermo Fisher Scientific, PA5-58716), anti-PMP70 (Abcam ab85550). The rat LCAD antibody was provided by Dr. Jerry Vockley at UPMC CHP and was previously characterized by us^10^. Ponceau S stain free total protein analysis is used as a reliable loading control^19^.

### Real-Time PCR

Real-time PCR analysis was performed as previously described to determine mRNA abundance^15^. cDNA was reverse-transcribed from 500 ng of total RNA with SuperScript First-Strand Synthesis System II (Thermo Fisher Scientific). Real-time PCR analysis was performed with gene specific primer oligos, SsoAdvanced SYBR Green Super-mix (Bio-Rad), and CFX96 Touch Real-Time PCR Detection System with C1000 Thermal Cycler (Bio-Rad). Cycling conditions were 95°C for 10 minutes, then 40 cycles of 95°C for 15 seconds and 60°C for 1 minute. Data were normalized to *Rn18S* as an endogenous control and analyzed using the 2^ΔΔCt^ method. Each primer sequence was shown in Table 1.

**Table 1.**
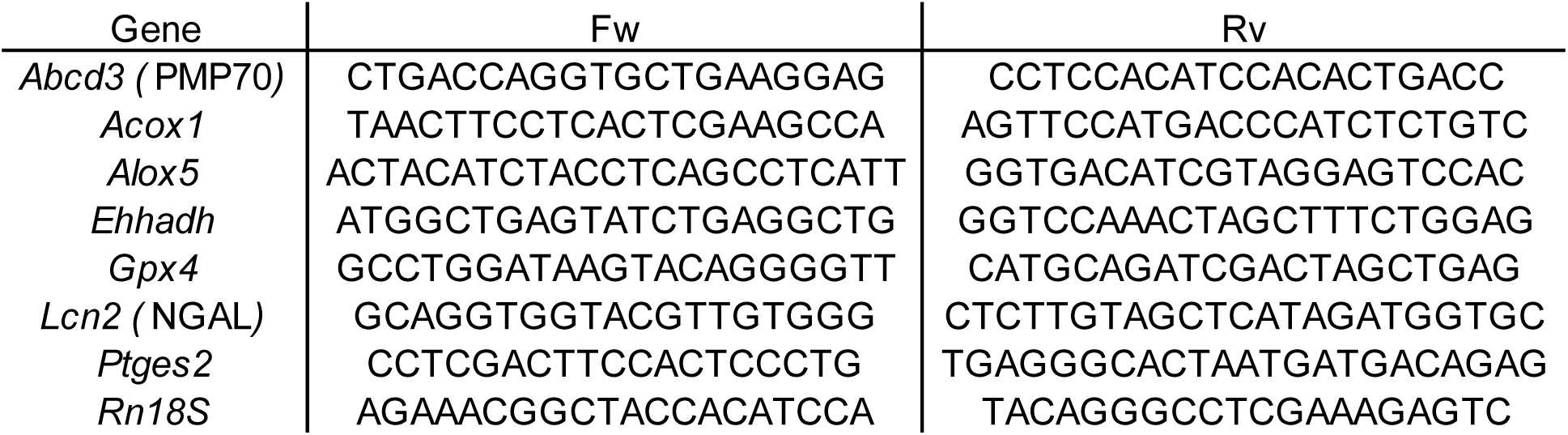
Primer sequences for qPCR.

### Tissue Section Analysis

Kidneys were fixed in 4% paraformaldehyde and embedded in OCT or paraffin. These tissues were sectioned at 10 μm or 4 μm, respectively. The kidney sections were stained with hematoxylin and eosin (H&E). H&E-stained slides were scored semi-quantitatively (0-4) for tubular damage in a blinded fashion with respect to tubular dilation, proteinaceous cast formation, and loss of brush border.

Immunostaining was performed as previously described^15^, with OCT- or paraffin-embedded tissues and with following primary antibodies: anti-LCAD (same as for Western blotting), anti-NGAL (R&D systems AF1857), anti-PEX5 (Cell Signaling Technology 830205), anti-PMP70 (Abcam ab85550), anti-SDHA (Cell Signaling Technology 11998), or LTL (Vector Laboratories FL-1321). Antibodies were used at 1:50 to 1:200, followed by conjugation with ALEXA^®^ fluorescent antibodies at 1:200.

TUNEL staining was performed by using the In Situ Cell Death Detection Kit (Roche) to evaluate cell death in paraffin embedded kidney tissue according to manufacturer instructions. TUNEL positive cells are quantified by ImageJ as previously described^20^.

### Electron transfer flavoprotein (ETF) fluorescence reduction assay

ETF fluorescence reduction assay analysis was performed to examine LCAD activity on FLUOstar Omega fluorescence spectrophotometer plate reader (BMG Labtech) as previously described^21^. Briefly, each 200 μl of assay reaction contained 50 μg kidney lysate, 25 μM palmitoyl-CoA. The reaction buffer contained 50 mM Tris-HCl, pH 8.0, 0.5% glucose, 2 μM recombinant pig ETF, 1.43 μl of glucose oxidase/ catalase mixture. The remaining activity of the KO mice should be from VLCAD (very long chain acyl-CoA dehydrogenase) which palmitoyl-CoA is also the favorable substrate.

### Radiolabeled Fatty Acid Oxidation Assays

^14^C-palmitate (PerkinElmer) was conjugated to BSA and used at 125 μM in 200 μl reactions as previously described^22^. ^14^CO_2_ was captured on filter papers soaked in 1M KOH and water-soluble ^14^C-labeled FAO products were separated by the method of Bligh and Dyer and subjected to scintillation counting^23^. Peroxisomal FAO was defined as the rate of palmitate oxidation in the presence of the irreversible mitochondrial FAO inhibitor etomoxir (100 μM). Data were normalized to protein concentration.

### Oroboros High-Resolution Respirometry

Freshly prepared kidney homogenates were analyzed with an Oroboros Oxygraph-2K using our previously published method^15^. Complex I respiration was defined as malate/ pyruvate/glutamate–driven oxygen consumption in the presence of ADP, whereas Complex II respiration was defined as succinate-driven oxygen consumption.

### Mitochondrial DNA (mtDNA) analysis

Total DNA was extracted using the DNeasy® Blood & Tissue kit (Qiagen 69504). mtDNA was quantified through a TaqMan-based assay as previously described^24–26^. Two sets of PCR primers, targeting the mitochondrial NADH dehydrogenase 1 (mt-ND1) gene and the histone deacetylase 1 (Hdac1) gene, were utilized to amplify the mtDNA and the nuclear DNA (nDNA), respectively^27^. PCR were performed on a CFX96 Touch Real-Time PCR Detection System (Bio-Rad, Inc.) under the following conditions: 95°C for 180 s; 45 cycles at 95°C for 15 s; and 60°C for 60 s. mtDNA copy number relative to the nDNA was calculated using the 2^-ΔΔCT^ method.

### F_2_-isoprostanes (IsoPs) and isofurans analysis

F_2_-IsoPs and isofurans were measured in the Vanderbilt University Eicosanoid Core Laboratory after lipid extraction from snap-frozen kidney samples using gas chromatography-mass spectrometry, as previously described^28^.

### Statistical Analyses

Data are presented as mean ± S.D. Prism 9.0.0 software (GraphPad) was used for statistical analysis. To determine whether sample data has been drawn from a normally distributed population, D’Agostino–Pearson omnibus test or Shapiro–Wilk test was performed. For parametric data, ANOVA with *post hoc* Tukey comparison was used for multiple group comparison and t test was used to compare two different groups. For nonparametric data, Mann–Whitney U test was used. The threshold of *p*<0.05 was set to consider data statistically significant.

## RESULTS

### LCAD lysine residue is hypersuccinylated in lysine deacylase sirtuin 5 knockout (*Sirt5^−/−^*) kidneys after ischemic AKI

We have previously shown that loss of the lysine deacylase sirtuin 5 confers renoprotection against AKI. We have shown that mitochondrial FAO enzymes are hypersuccinylated to inhibit their activity in *Sirt5^−/−^*kidneys. Metabolic adaptation to blocked mitochondrial FAO involves compensatory FAO in the peroxisome, resulting in mitigation of oxidative stress and renoprotection^15^. We now show that succinylation of LCAD residue K322 is increased by 16-fold in *Sirt5^−/−^* kidney post-AKI (Figure 1). Succinylation of residue K322, which is near the LCAD active site, suppresses enzymatic activity^29^. Because LCAD can be a source of mitochondrial H_2_O_2_, we hypothesized that the loss-of-function of LCAD attenuates oxidative stress to protect against AKI^10^. We confirmed that WT mice express LCAD in the kidney including LTL^+^ PTECs whereas *LCAD^−/−^*kidneys do not (Figure 2A-B). The faint band present in *LCAD^−/−^* kidney tissue is due to slight cross-reactivity of the LCAD antibody to other acyl-CoA dehydrogenases of the same size (unpublished). No difference was noted in renal function and renal histology in the absence of injury between *LCAD^−/−^*and WT kidneys.

**Figure 1.**
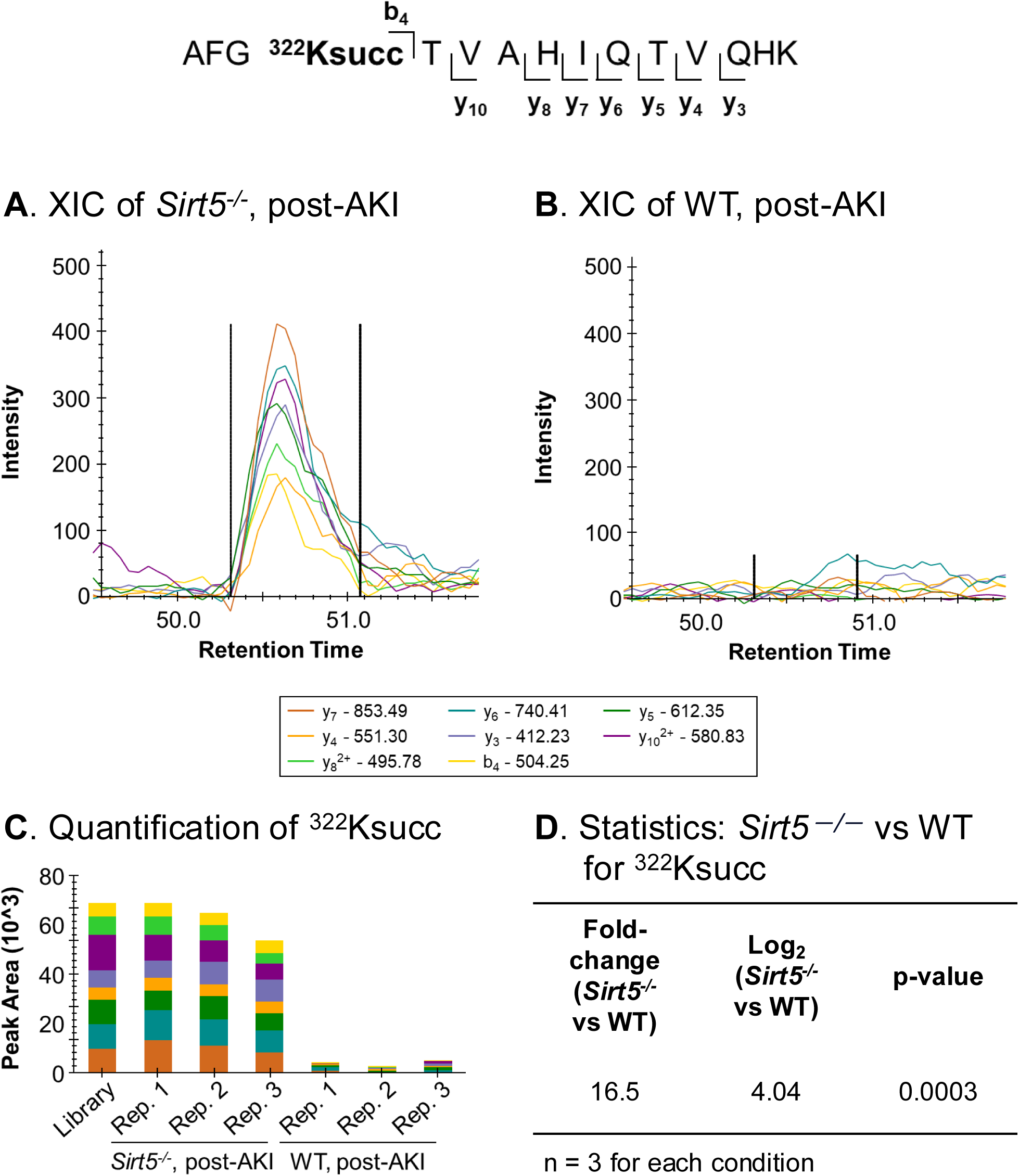
Mass spectrometry revealed hypersuccinylation of lysine K322 of LCAD in *Sirt5−/−* kidneys post-AKI. Extracted ion chromatograms (XICs) of the peptide AFG**322Ksucc**TVAHIQTVQHK (precursor ion at m/z 441.99, z = 4+) of LCAD in kidney tissues from a (A) a *Sirt5 −/−* biological replicate and a (B) WT biological replicate, post-AKI. (C) Quantification of the succinylated peptide in the three *Sirt5 −/−* replicates and the three WT replicates, showing peak areas as determined in Skyline. (D) Statistical analysis confirmed the increased succinylation level of K322 of LCAD in *Sirt5-/-* vs WT kidneys, post-AKI.

**Figure 2.**
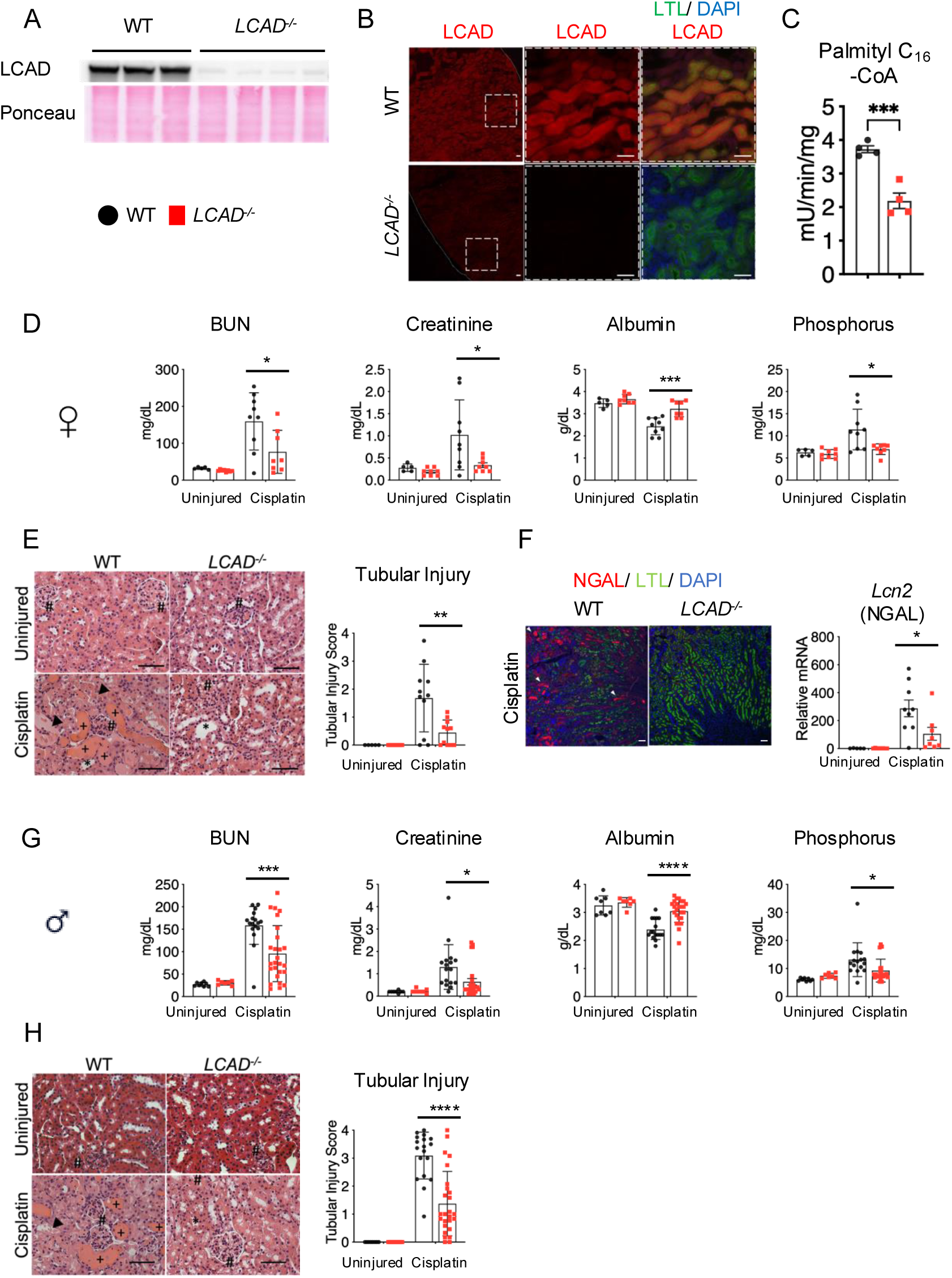
*LCAD−/−* kidneys are protective against cisplatin-AKI in sex-independent manner. (A) Anti-LCAD immunoblot in *LCAD−/−* versus WT mouse kidney lysates (B) Anti-LCAD immunostaining in *LCAD−/−* versus WT mouse kidney, co-stained with Lotus Tetragonolobus Lectin (LTL) (C) *LCAD−/−* versus WT mouse kidney homogenates were tested for palmitoyl-CoA (C16-CoA) dehydrogenase activities. n=3-4, (D-F) female mouse studies (G-H) male studies (D or G) Serum analyses for BUN, creatinine, Albumin and Phosphorus indicate that the level of renal functional impairment is decreased in *LCAD−/−* mice 3 days post cisplatin treatment in sex independent manner. (E or H) Hematoxylin and eosin-stained kidney tissues demonstrate that renal tubular injury is mitigated in *LCAD−/−* kidneys 3 days post cisplatin treatment in sex independent manner. Glomeruli (#), proteinaceous cast (+), dilated tubule (*) n=8-11. Representative images and semiquantitative scoring for renal tubular damage at day 3 post cisplatin treatment are shown. (F) *LCAD−/−* kidneys decreased mRNA (*Lcn2*) and immunostained tubular cells for Neutrophil gelatinase-associated lipocalin (NGAL) in whole kidneys 3 days after cisplatin treatment. Scale bars: 50µm.\**p*<0.05; \*\*\**p*<0.001; \*\*\*\**p*<0.0001. One-way ANOVA *post hoc* Tukey multiple comparison

Electron transfer flavoprotein (ETF) fluorescence reduction assay was used to measure acyl-CoA dehydrogenase enzyme activity. Mitochondria isolated from WT or *LCAD^−/−^* kidneys were compared using palmitoyl (C_16_)-CoA as a substrate, representing the combined activities of LCAD, very long-chain acyl-CoA dehydrogenase (VLCAD), and ACAD9 (mostly LCAD and VLCAD). Interestingly, *LCAD^−/−^* kidney mitochondria show only ≈40% reduction of combined acyl-CoA dehydrogenase activity with C_16_-CoA (Figure 2C), whereas ∼65% reduction was reported in *LCAD^−/−^* liver mitochondria^30^. This suggests that, in kidneys, VLCAD plays a significant role in mitochondrial long-chain FAO.

### *LCAD^−/−^* kidneys are protected against distinct AKI models by cisplatin or renal IRI in mice

We next treated single high dose of cisplatin (20 mg/kg b.w. i.p.) to induce nephrotoxic AKI on WT or *LCAD^−/−^* female mice. *LCAD^−/−^* mice had decreased renal injury 3 days post cisplatin, assessed by decreased serum levels of BUN, creatinine, and phosphorus, with an attenuated reduction in serum albumin (Figure 2D). Low serum albumin levels are known risk factors for human cisplatin AKI^31^. We have previously observed that serum albumin is significantly elevated in *LCAD^−/−^* lavage fluid^32^. Histology and immunostaining further confirmed reduced renal tubular injury as well as reduced expression of NGAL (a kidney injury marker) in *LCAD^−/−^*kidney tissue (Figure 2E-F). We also performed the corresponding experiments in male mice and observed consistent results with the females, suggesting that our findings are sex independent (Figure 2G-H).

*LCAD^−/−^* and WT male mice were also subjected to an ischemic AKI model in which using unilateral renal ischemia/reperfusion injury followed by delayed contralateral nephrectomy 1 day before the samples were collected on day 7 (Figure 3). We obtained consistent results from the cisplatin-AKI model, suggesting that *LCAD^−/−^* kidneys are protective against multiple distinct AKI models.

**Figure 3.**
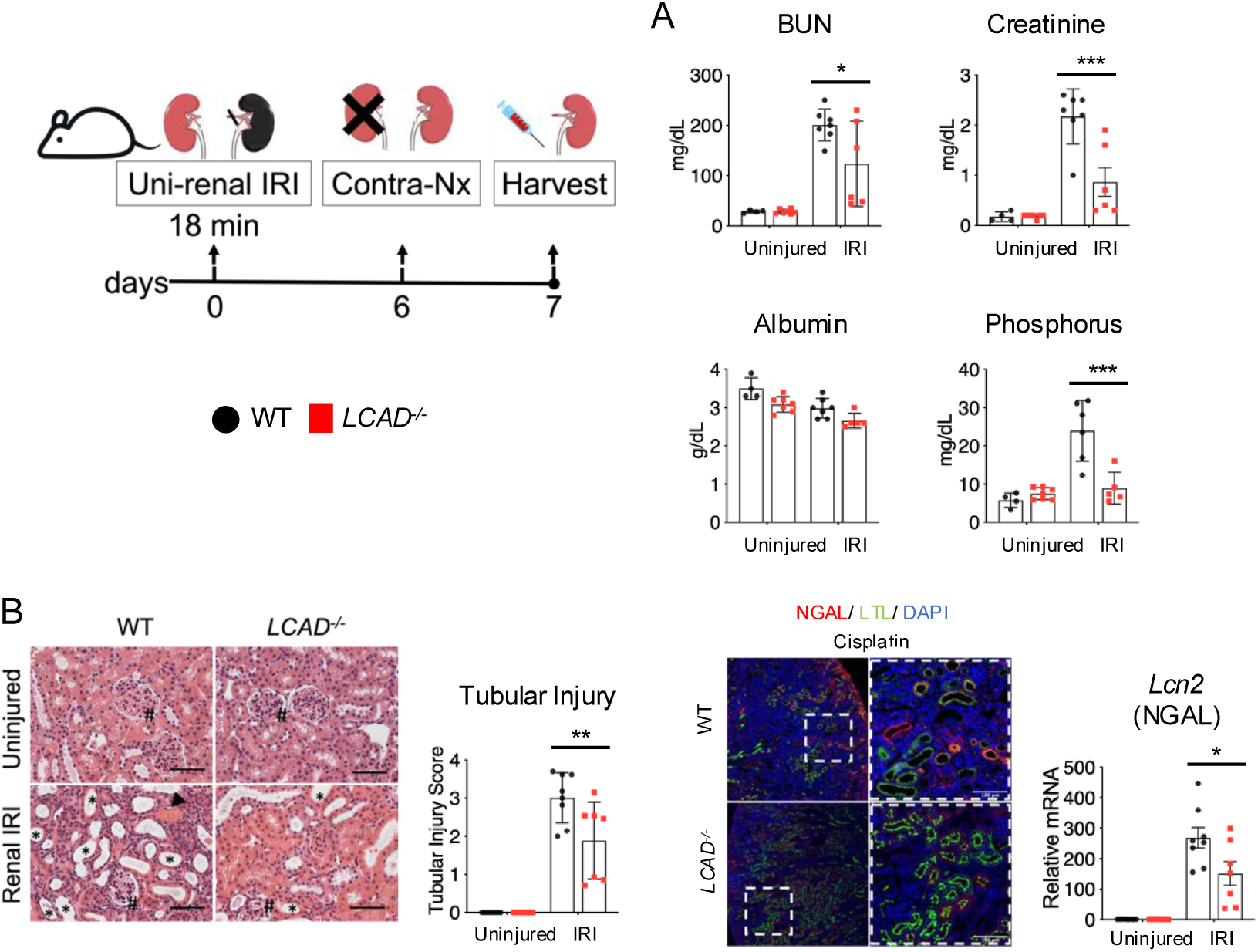
*LCAD−/−* kidneys are protective against ischemic-AKI. (A) Serum analyses for BUN, creatinine, Albumin and Phosphorus indicate that the level of renal functional impairment is decreased in *LCAD−/−* mice 7 days post renal IRI in male mice. (B) H&E-stained kidney tissues demonstrate that renal tubular injury is mitigated in *LCAD−/−* kidneys 7 days postrenal IRI. Glomeruli (#), proteinaceous cast (+), dilated tubule (*) n=6-7. Representative images and semiquantitative scoring for renal tubular damage at day 7 post renal IRI are shown. *LCAD−/−* kidneys decreased mRNA (*Lcn2*) and immunostained tubular cells for Neutrophil gelatinase-associated lipocalin (NGAL) in whole kidneys 3 days after cisplatin treatment. Scale bars: 50µm.\**p*<0.05; \*\**p*<0.01; \*\*\**p*<0.001. One-way ANOVA *post hoc* Tukey multiple comparison

### Mitochondrial function and peroxisomal FAO are preserved in *LCAD^−/−^* kidneys after cisplatin-AKI

We analyzed mitochondrial respiration on carbohydrate-derived substrates using the Oroboros Oxygraph-2K high-resolution respirometer with *LCAD^−/−^*or WT kidneys ± cisplatin-AKI. As in liver, *LCAD^−/−^* mitochondria in uninjured kidneys showed no change in respiratory chain function^10^. Interestingly, respiratory chain function is also preserved in injured *LCAD^−/−^* mitochondria while injured WT mitochondria demonstrated a significant decrease in respiration on the Complex I substrate pyruvate (Figure 4A). Succinate dehydrogenase complex flavoprotein subunit A (SDHA) is a component for SDH, which is a critical mitochondrial enzyme for the TCA cycle and the electron transport chain. Consistent with the Oroboros respirometric data, mitochondrial (mt) DNA level is increased and SDHA expression is preserved in injured *LCAD^−/−^* kidneys (Figure 4B-D).

**Figure 4.**
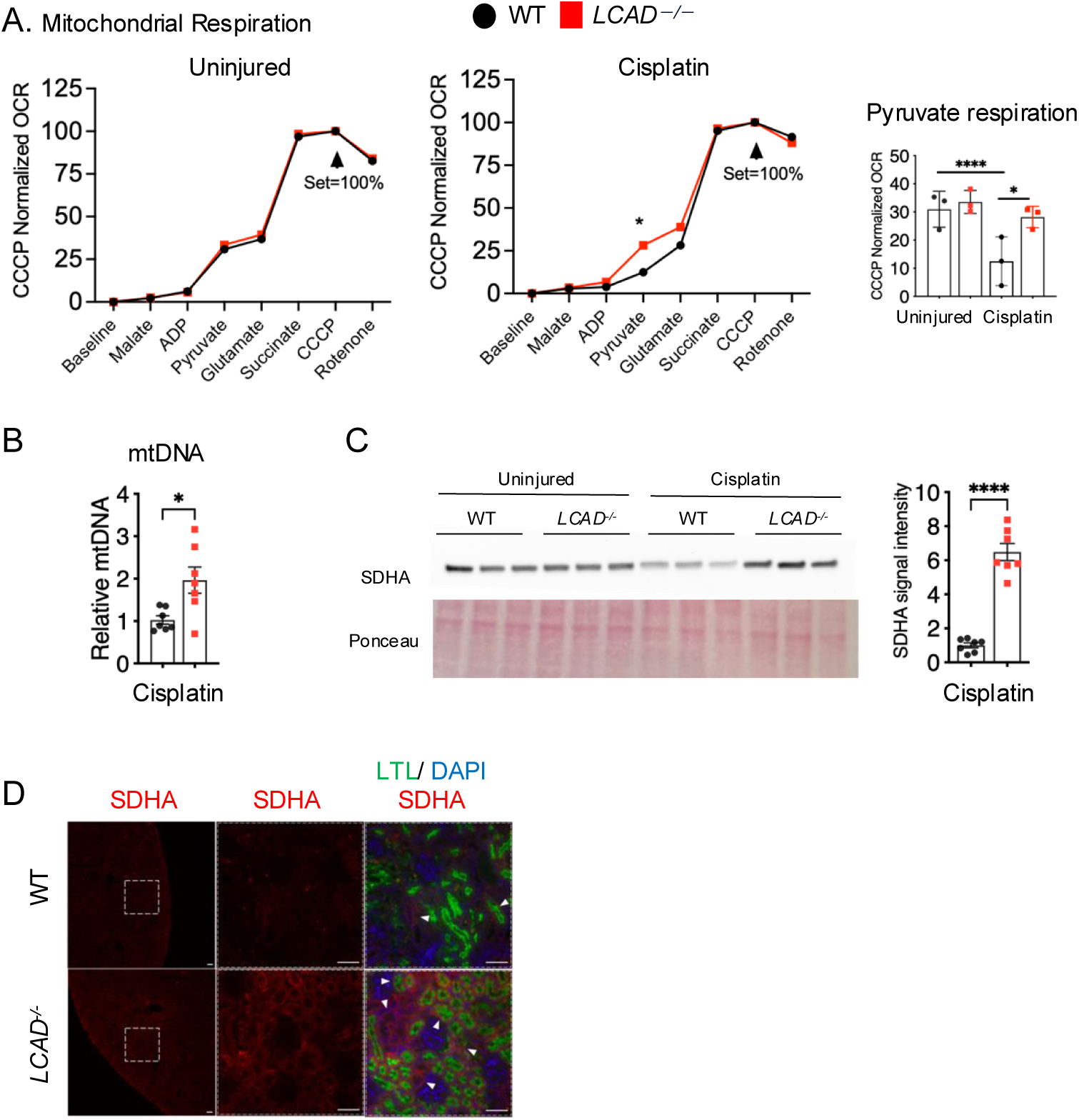
*LCAD−/−* kidneys preserve mitochondrial function in cisplatin-AKI. (A) Oroboros O2k respirometry revealed that Complex I of the electron transport chain was increased in *LCAD-/-* kidneys compared with WT 3 days after cisplatin treatment. (B) Mitochondrial (mt) DNA levels are increased in *LCAD−/−* kidneys (C) *LCAD−/−* kidneys increased immunoblotted succinate dehydrogenase complex flavoprotein subunit A (SDHA) expression in whole kidney lysates 3 days after cisplatin treatment. A representative blot image and the quantified data for cisplatin treated samples are shown (D) Immunostained SDHA in whole kidneys of WT versus *LCAD−/−* 3 days after cisplatin treatment. Scale bars: 50µm.\**p*<0.05; \*\*\*\**p*<0.0001. One-way ANOVA *post hoc* Tukey multiple comparison or *t*-test

We and others have previously shown that enhanced peroxisomal function is linked to renoprotection against AKI^15,18,33,34^. We therefore interrogated *LCAD^−/−^* kidneys for peroxisomal FAO after cisplatin AKI. The catabolism of a radiolabeled long-chain fatty acid, ^14^C-palmitate, was followed to ^14^CO_2_ and water-soluble short-chain fatty acids in kidney homogenates in the presence of etomoxir (an irreversible inhibitor of mitochondrial FAO). Etomoxir-resistant FAO is ascribed to peroxisomes. While there is no change between uninjured WT and *LCAD^−/−^* kidney, injured *LCAD^−/−^* kidney displayed dramatically higher peroxisomal FAO after cisplatin-AKI (Figure 5A). It is consistent that two peroxisomal FAO genes (*Acox1* and *Ehhadh*) were upregulated in injured *LCAD^−/−^* kidneys (Figure 5B). *LCAD^−/−^* kidneys were confirmed to have greater abundance of peroxisomal membrane markers PMP70 and PEX5 in the kidneys after cisplatin-AKI suggesting preservation not only of function but also the number of peroxisomes (Figure 5C-D).

**Figure 5.**
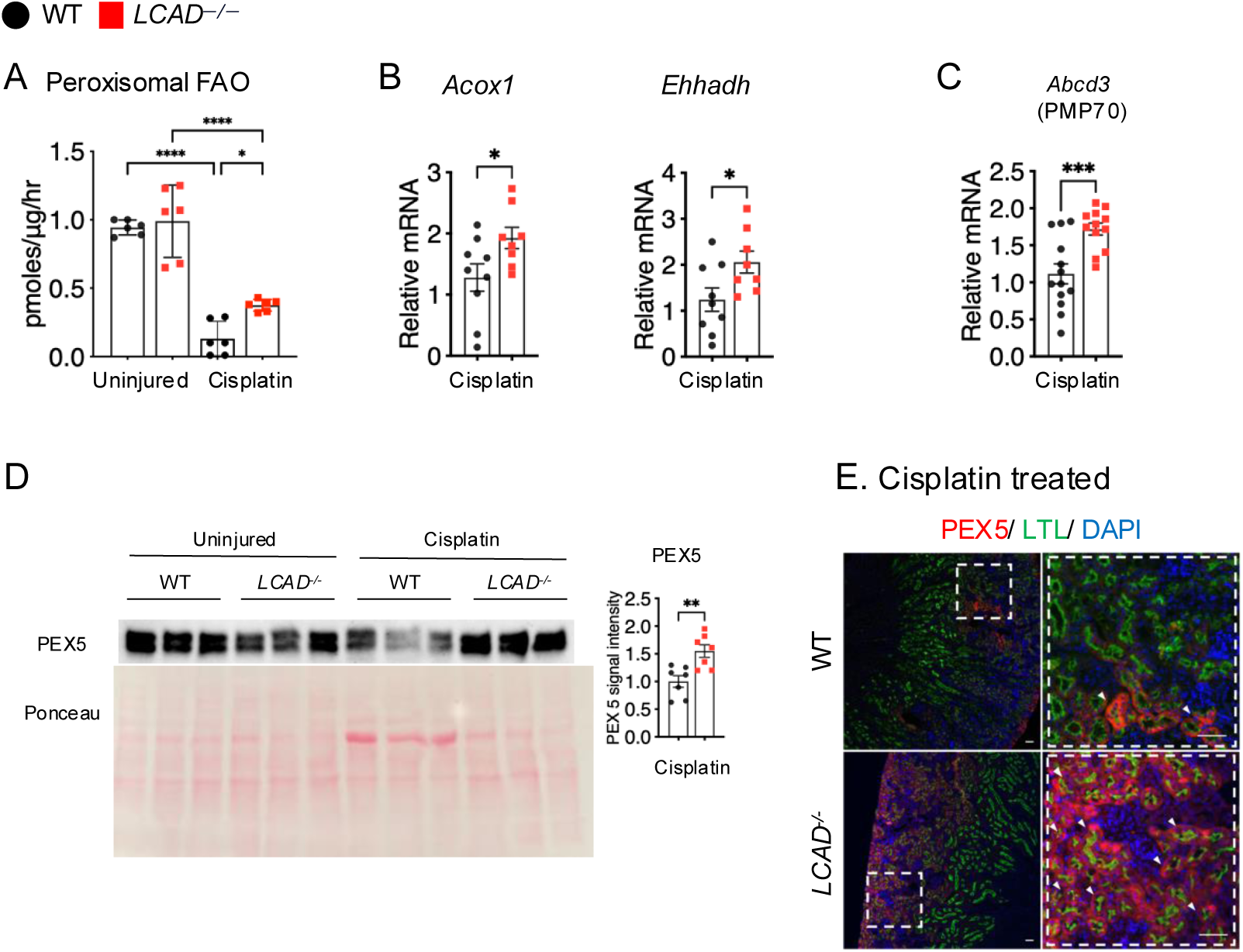
*LCAD−/−* kidneys enhances peroxisomal fatty acid oxidation (FAO) in cisplatin-AKI. (A) Etomoxir-insensitive 14C-palmitate oxidation, which represents peroxisomal FAO, was decreased in kidney homogenates 3 days after cisplatin treatment in WT. *LCAD−/−* kidneys increase peroxisomal FAO after 3 days cisplatin treatment as compared to WT. (B) *LCAD−/−* kidneys increases mRNA levels of peroxisomal FAO associated genes (*Acox1* and *Ehhadh*). (C) mRNA levels of PMP70 (peroxisomal marker) is increased in *LCAD−/−* kidneys 3 days after cisplatin treatment. (D) Immunoblotted PEX5 (peroxisomal marker) is increased in *LCAD−/−* kidneys 3 days after cisplatin treatment. (E) PEX5 immunostaining images are also shown. Scale bars: 50µm.\**p*<0.05; \*\**p*<0.01, \*\*\**p*<0.001, \*\*\*\**p*<0.0001. One-way ANOVA *post hoc* Tukey multiple comparison or *t*-test

### Oxidative stress and ferroptotic cell death are decreased in *LCAD^−/−^*kidneys after cisplatin-AKI

Consistent with our central hypothesis, an *in vivo* marker of oxidative stress, isofuran/F_2_-isoprostane^28^, was decreased in the *LCAD^−/−^* kidneys after cisplatin AKI (Figure 6A). Ferroptosis is a regulated form of cell death by iron-dependent lipid peroxidation in which mitochondrial oxidative stress plays an important role^35^. Abnormal fatty acid metabolism is essential for the delivery of the molecular signals to induce ferroptotic cell death^36^. We performed TUNEL analysis, a universal assay for programmed cell death including ferroptosis, and showed that *LCAD^−/−^* kidneys had decreased TUNEL^+^ cells after cisplatin AKI (Figure 6B). We further tested three ferroptosis markers Gpx4 (down-regulated in ferroptosis), Alox5 and Ptges2 (upregulated in ferroptosis). We have shown that *LCAD^−/−^* kidneys have increased levels of Gpx4 and decreased levels of Alox5 and Ptges2, which suggests reduced ferroptosis (Figure 6C-D).

**Figure 6.**
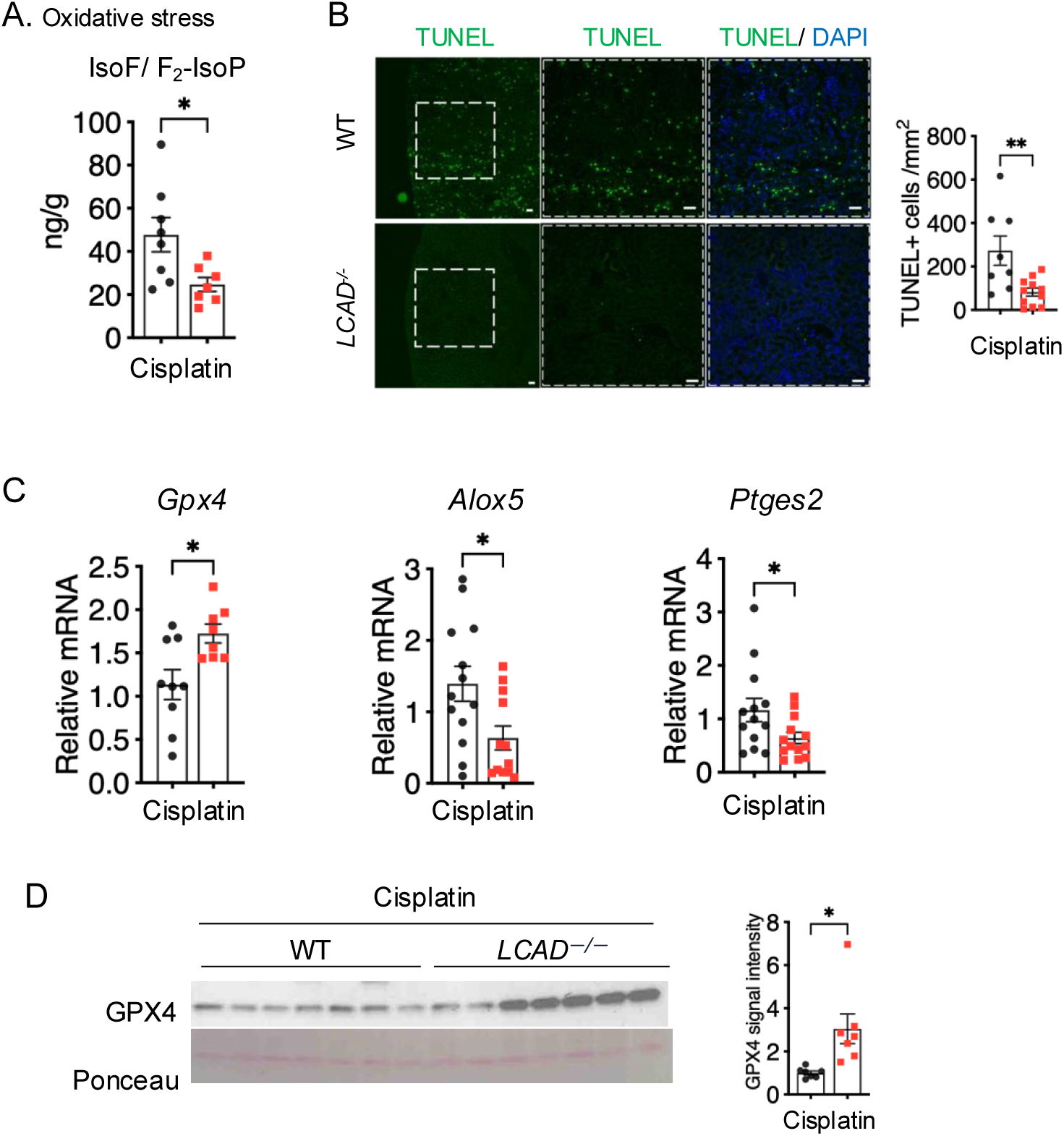
*LCAD—/—* kidneys mitigate oxidative stress and ferroptosis in cisplatin-AKI. (A) The *in vivo* oxidative stress marker, isofuran/F2-Isoprostane (IsoF/F2-IsoP) ratio, is decreased in *LCAD-/-* kidneys 3 days after cisplatin treatment. (B) TUNEL-positive dying cells are decreased in *LCAD-/-* kidneys 3 days after cisplatin treatment. (C) Ferroptosis associated genes (Alox5 and Ptgs2) are decreased while Gpx4 is increased in *LCAD-/-* kidneys 3 days after cisplatin treatment. (D) Immunoblotted GPX4 expression is increased in *LCAD-/-* kidneys 3 days after cisplatin treatment. Scale bars: 50µm.\**p*<0.05. *t*-test

## DISCUSSION

Proximal tubules, which are the major site of injury during AKI, are rich in mitochondria and rely on FAO for their high energy demands^4,5^. During AKI, mitochondria are damaged, FAO is compromised, and toxic lipids accumulate, thereby leading to oxidative stress^3,37–40^. Thus, it has been largely held that dysfunctional mitochondrial FAO promotes AKI and that augmenting mitochondrial FAO will protect against injury and accelerate recovery^41–43^. Agents that indirectly stimulate mitochondrial function and FAO, such as PGC1α overexpression^44^, the AMPK agonist AICAR, PPARα agonist drugs, and a compound called C75 that is believed to activate a critical FAO enzyme known as carnitine palmitoyltransferase-I (CPT1) have all been associated with protection against AKI^45–47^. At the same time, etomoxir, an irreversible inhibitor of CPT1 that strongly *inhibits* mitochondrial FAO, also protects against AKI^48^. Thus, the role of mitochondrial FAO in protection vs. exacerbation of renal injury in AKI is unclear.

These apparently discrepant data regarding mitochondrial FAO during AKI might be reconciled by considering peroxisomal FAO. Proximal tubules are also rich in peroxisomes, which possess a biochemically parallel FAO pathway to that of the mitochondria that does not directly produce ATP. While PPARα agonists increase expression of mitochondrial FAO genes, their effects on the peroxisomal FAO gene expression are much greater, at least in rodents^49^. Agents that have been shown to protect against AKI such as AICAR and C75 all activate PPARα and will thus increase peroxisomal biogenesis and FAO^48,50–54^. PGC1a has also been shown to increase peroxisomal abundance and activity^55^. There appears to be considerable crosstalk and cooperation between the peroxisomal and mitochondrial FAO pathways. Some substrates are partially chain-shortened in peroxisomes and then passed on to mitochondria^56^. Further, inhibiting flux through mitochondrial FAO causes a compensatory increase in peroxisomal FAO and vice versa^57,58^.

Mitochondrial ROS is well known to be a driver of numerous disease states in the kidney, and FAO is the largest contributor to this ROS. In one study of diabetic kidney, the pathological increase in ROS was ascribed completely to FAO^3^. Importantly, the source of this ROS was found to be upstream of the electron transport chain. The leaked electrons were coming either directly from the acyl-CoA dehydrogenases, which catalyze the first step in β-oxidation, or from their redox partners electron transferring flavoprotein (ETF) and ETF dehydrogenase. Others have shown that ETF and ETF dehydrogenase can leak electrons^59,60^, and we have shown that amongst the acyl-CoA dehydrogenases, it is LCAD that is most susceptible to electron leakage to oxygen. LCAD has a wider substrate-binding pocket than other acyl-CoA dehydrogenase family members, which allows oxygenated solvent partial access to the electrons. The result is electrons jumping to oxygen and direct formation of H_2_O_2_. While LCAD downregulation is observed in many cancers to reduce H_2_O_2_ levels^11,12^, re-expression of LCAD in liver cancer cell lines by stable transfection increases ROS^12^.

Here, we showed that partial suppression of mitochondrial FAO by ablation of LCAD was sufficient to reduce the severity of tissue injury induced by two forms of AKI. The mechanism of protection may involve a reduction in mitochondrial ROS coincident with increased flux of fatty acids through the peroxisomal FAO pathway. The peroxisomal FAO pathway also produces H_2_O_2_, but peroxisomal catalase reverts the H_2_O_2_ back to O_2_ and water. The action of catalase, which in essence regenerates oxygen, allows FAO to proceed with a reduced oxygen cost which could be particularly beneficial during ischemic AKI. One weakness of our study that should be noted is that we did not directly measure peroxisomal FAO flux, but rather, used the mitochondrial inhibitor etomoxir to differentiate between the two pathways, and etomoxir has been suggested to have off-target effects on the electron transport chain and on triglyceride synthesis^61–63^. However, these off-target effects were all reported in experiments where etomoxir was applied to cultured cells for extended periods of time. Here, we measured FAO ± etomoxir over a 1 hr interval using tissue homogenates. Further, any artifacts introduced by off-target effects would be equal across all of the assays. Regardless, future studies are needed to more accurately quantify the degree of peroxisomal FAO induction in proximal tubules upon ablation of the mitochondrial pathway.

Our studies here implicate suppression of LCAD as a contributor to the renoprotection that we previously reported in *Sirt5^−/−^* mice^15^. In that study, we noted hypersuccinylation of enzymes representing virtually every step of mitochondrial FAO, including LCAD K322. Here, we chose to study LCAD due to previous work demonstrating a direct role of K322 in LCAD enzyme activity^29^. *LCAD^−/−^* mice largely phenocopied the renoprotection seen in Sirt5 mice, with similar degrees of injury mitigation. While it is tempting to interpret our data as suggesting that LCAD suppression explains the benefits of Sirt5 inhibition, we did not directly interrogate a Sirt5-LCAD axis. Inhibition of mitochondrial FAO at other steps in the pathway may very well produce similar results, which is a topic of future investigation. Translationally, LCAD may be an attractive target for AKI intervention. In rodents LCAD is broadly expressed and plays a critical role in tissues where mitochondrial FAO is indispensable for energy production, such as heart and muscle. In humans, however, LCAD expression is restricted to kidney, liver, lung, and pancreas^10^, such that inhibition of LCAD in humans would not compromise heart and muscle as it would in rodents. Studies are underway to determine the optimal means by which to exploit this novel mechanism for the prevention and treatment of AKI.

## Supporting information

Unedited blots and gels

## FUNDING

This work was supported by NIDDK R01-DK121758 (SSL), NIDDK R01-DC090242 (ESG), NICHD R01-HD103602 (ESG), NIDDK K01-DK133635 (TC), UPMC CHP RAC Start-up grant (TC), Samuel and Emma Winters Foundation grant (TC), UPMC VMI P3HVB (TC), Fukushima Medical University Postdoctoral Fellowship (AO), Richard K. Mellon Postdoctoral Fellowship (KEP) , and NIH-shared instrumentation grant 1S10 OD016281 for the Triple TOF 6600 (Buck Institute).

## AUTHOR CONTRIBUTIONS

TC and AO designed and performed experiments, provided intellectual input, and wrote the manuscript. YZ, JB, SSB, KEP, BBZ, and ACR performed experiments and provided intellectual input. BS, ESG and SSL designed and performed experiments, edited the manuscript, provided intellectual input, and oversaw the project. All authors approved the final version of this manuscript.

## ACKNOWLEDGEMENTS

The University of Pittsburgh Center for Biologic Imaging provided microscopy support. We thank the support of the UPMC CHP Histology Core Laboratory for histology assistance, of the Kansas State University Veterinary Diagnostic Laboratories for serum chemistry, and of the Vanderbilt Eicosanoid Core Laboratory for isofuran and F_2_-Isoprostane analysis.

## CONFLICT OF INTEREST

None

## DATA SHARING STATEMET

The data supporting the findings of this study are publicly available through Mass Spectrometry Interactive Virtual Environment (MassIVE; ftp://massive.ucsd.edu/v02/MSV000083439/) with MassIVE ID: MSV000083439; ProteomeXchange ID: PXD012696.

## Notes

### Competing Interest Statement

The authors have declared no competing interest.

