## Supplementary material for "Loss of long-chain acyl-CoA dehydrogenase protects against acute kidney injury": Unedited blots and gels

### Uncropped Western blot images

#### Figure 1A (LCAD)

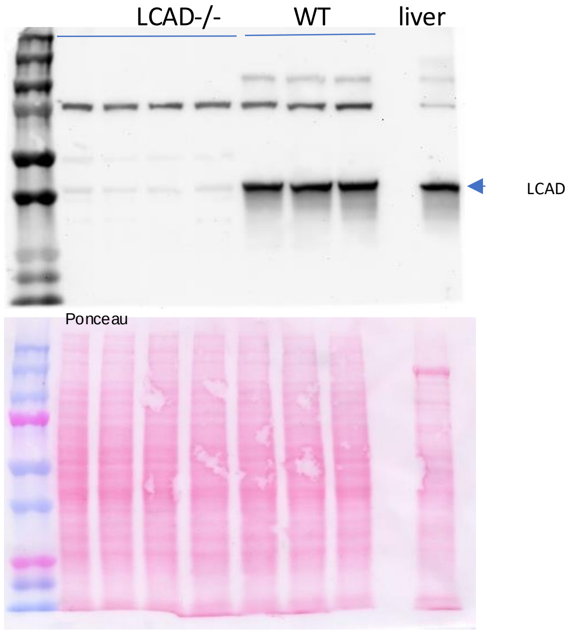


*Figure 1A image has been horizontally inverted to show WT samples on the left and LCAD KO samples on the right.

#### Figure 4C (SDHA)

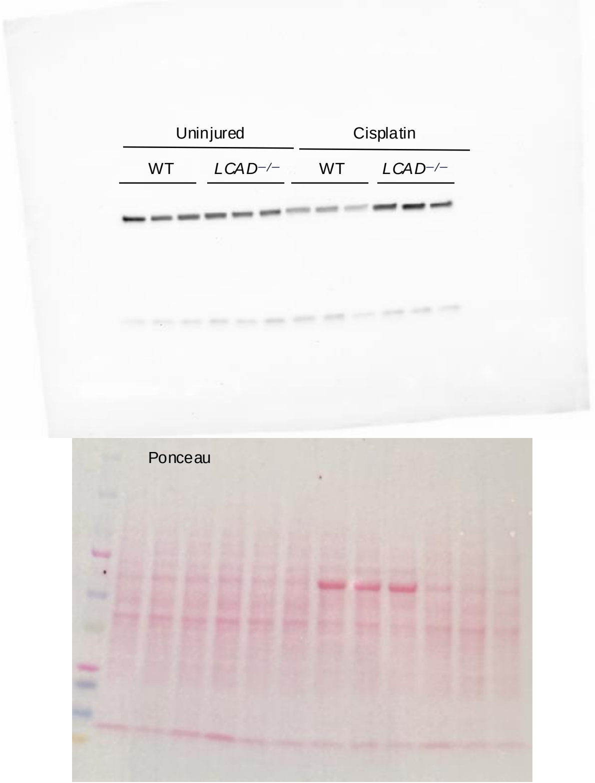


#### Figure 5D (PEX5)

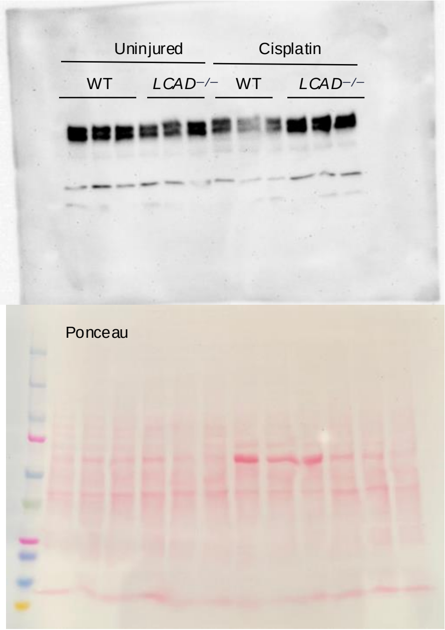


#### Figure 6D (GPX4)

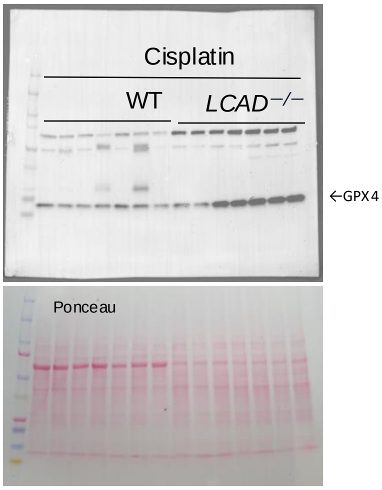
